# Exogenous Amyloid Sequences: Their Role in Amyloid-Beta Heterotypic Aggregation

**DOI:** 10.1101/2025.01.24.634659

**Authors:** Jofre Seira Curto, Marina Romero Ruiz, Genís Pérez Collell, Sandra Villegas, Maria Rosario Fernandez, Natalia Sanchez de Groot

## Abstract

Protein aggregation is a complex process influenced by environmental conditions and interactions between multiple molecules, including those of exogenous origin. Although *in vitro* simulations of aggregation are crucial for advancing research, few studies explore cross-seeding as a repeating event, despite the potential for such events when proteins circulate through the body. Here, we investigated the impact of exogenous amyloid sequences derived from the gut microbiota on the heterotypic aggregation of Aβ peptides. We utilized ten 21-amino acid peptides derived from bacterial genomes, previously shown to interfere with Aβ40 aggregation and induce memory loss in *Caenorhabditis elegans*. Through consecutive cross-seeding assays with Aβ40 and Aβ42, we analyzed the effects of these peptides on aggregation kinetics and seed propagation. Our findings indicate that exogenous molecules can influence Aβ’s aggregation process, altering the fibrils’ properties. Based on this, we introduce the “Interaction History” concept, where prior interactions shape the aggregation and propagation of Aβ peptides. This work supports the idea that environmental factors, such as microbial amyloids, can contribute to the heterogeneity and progression of amyloid-related diseases. Our results highlight the need for therapeutic strategies targeting diverse amyloid configurations and reinforce the importance of considering exogenous sequences as additional triggers in AD pathology.

## Introduction

Protein aggregation is associated with many neurodegenerative diseases. In particular, the formation of amyloid plaques has been widely described in the brains of Alzheimer’s disease (AD) patients[1–5]. These deposits are composed principally of amyloid beta peptide (Aβ), which exist in two major isoforms: Aβ40, the most abundant in the body, and Aβ42, the main component of amyloid plaques [4,6] (Figure 1A). These two isoforms, differ in length and aggregation propensity and can interact with each other, producing different outcomes[2,7– 11]. For instance, *in vivo* studies have shown that Aβ40 overexpression inhibits Aβ42 deposition, suggesting a potential protective role for Aβ40 [9,12,13]. The balance between Aβ40 and Aβ42 is influenced by isoforms of the apolipoprotein E gene (APOE). Among these, APOE4 is particularly significant, as it promotes Aβ42 formation, exacerbating aggregation and accelerating disease progression[5,14]. At the molecular level, interlaced amyloid fibrils containing both Aβ40 and Aβ42 have been observed, supporting that these isoforms can interact during the aggregation process [9]. Similarly, compositional analyses of amyloid plaques extracted from patients’ brains also revealed heterogeneous aggregation events[15,16].

Beyond the brain, Aβ aggregation can also involve systemic interactions, and increasing evidence supports the idea that Aβ peptides can cross-seed and propagate aggregation across different regions of the body, potentially contributing to the spread of amyloid pathology[17] (Figure 1A). Aβ is present in both the central nervous system (CNS) and the peripheral circulation, maintained by a dynamic equilibrium between these two pools[18]. However, disruptions in this balance can have significant consequences for Aβ homeostasis. While circulating Aβ can enter the brain, elevated peripheral Aβ levels may impair its clearance from the CNS, leading to increased accumulation[18,19]. This can occur when blood-brain barrier (BBB) is compromised, allowing Aβ peptides to infiltrate into the brain and interact with neurons[5], potentially influencing aggregation dynamics. Additionally, circulating macrophages[20] and blood monocytes[19] may also serve as alternative transport mechanisms, facilitating the movement of Aβ seeds from the periphery into the brain.

Notably, Aβ aggregation is not restricted to interactions between different isoforms[8,21]; but it is also influenced by its interaction with other proteins, such as amylin[22] or alpha-synuclein[23–25], as well as non-protein molecules like RNA[26,27] and metals[28–30]. These other molecules, along with changes in local environmental conditions (e.g., pH), can produce amyloid aggregates with distinct structural and biophysical properties [31–35]. Beyond the diversity induced by external factors, amyloid fibril polymorphism can also emerge withing a single aggregation process, with multiple structures populating the different aggregational phases, and even after reaching the plateau[34,35]. Although lower fibril diversity has been observed *in vivo*, structurally distinct amyloid fibrils, also known as strains, have been linked to different clinical and pathological outcomes in AD[36,37]. In this line, it has been observed a connection between environmental factors and lifestyle with AD development[38], highlighting the potential impact of exogenous molecules in this process.

AD’s research is gradually expanding to include the microbiota (Figure 1A), which plays a significant role in influencing the host’s immune and regulatory processes[39]. In this context, microbiome-derived proteins can be exported and digested, for example, in the gut, resulting in molecules capable of influencing the aggregation of host proteins[39,40]. This aligns with the reported potential of prion-like proteins for transcellular propagation[17] and interspecies transmission[25,41–44]. An increasing body of evidence suggests that microbial cells and their products can circulate through the host body, potentially reaching the brain via the vagus nerve or bloodstream[39,45], thereby able to influencing aggregation processes far away from the original site of infection (Figure 1A).

Overall, rather than a static process, amyloid aggregation is iterative, dynamic, and environment-dependent[22,23,35,24–30,34], leading to an energy landscape with multiple local minima, each corresponding to distinct fibrillar polymorphs[34,35,46]. Consequently, the introduction of an exogenous fibrillar seed can perturb this landscape by stabilizing specific polymorphs, shifting aggregation pathways, or promoting the formation of novel aggregate structures. In this context, we investigated how exogenous proteins, which may be present in the body, influence Aβ aggregation dynamics. To explore this, we used a collection of prion-like sequences derived from the gut microbiome[47] (Figure 1B), which have recently been shown to interfere with Aβ40 aggregation *in vitro* and induce memory loss when ingested by *C. elegans*.

**Figure 1.**
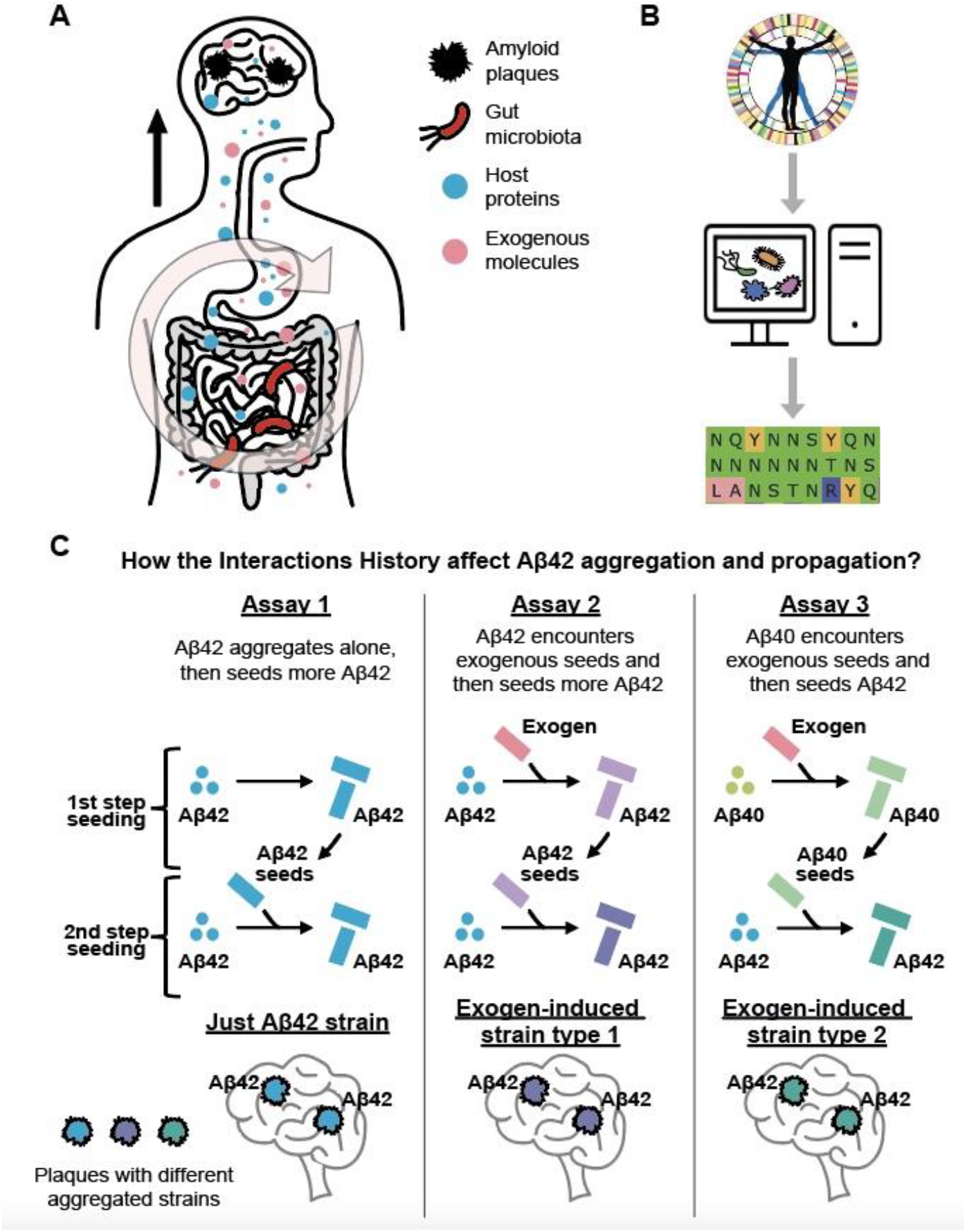
Exogenous prion-like sequences can affect Aβ aggregation and propagation. A) Microbial cells and their products can circulate through the body, encountering host proteins, reaching the brain, and influencing aggregation processes. B) Computational analysis of the human microbiome has shown the presence of several genes coding for prion-like proteins[47]. C) Schematic representation of the experimental approach used to study the effect of Interaction History on Aβ42 aggregation and propagation. We consider three hypothetical scenarios where two consecutive seeding steps influence Aβ42 aggregation. Scenario 1: Only Aβ42 molecules act as monomers and seeds in both seeding steps. Scenario 2: Aβ42 aggregates in the presence of an exogenous molecule, and subsequently, these aggregates seed new Aβ42 monomers. Scenario 3: Aβ40 aggregates in the presence of an exogenous molecule, and subsequently, these Aβ40 aggregates seed Aβ42 monomers. Our results suggest that if these three different aggregation pathways occur in the body, they may lead to the formation of different aggregated Aβ42 strains. Dots represent monomeric proteins, while bars indicate aggregated structures. Colors denote different aggregate identities: Blue for unexposed Aβ42, red for exogenous amyloids, yellow for unexposed Aβ40, light green for aggregate Aβ40, and the other colors represent different aggregated Aβ42 polymorphs.

In the present study, we aim to investigate how these bacterial sequences influence both the formation of Aβ42 aggregates and, more importantly, the seeding properties of these aggregates, particularly their ability to trigger subsequent aggregation events. To this end, we focus on the cross-interactivity between Aβ40 and Aβ42[9] (Figure 1C). We observed that incubating Aβ with bacterial sequences accelerated aggregation and produced deposits with varying seeding capacities (Figures 2-4). These findings suggest that the introduction of additional molecular factors may contribute to amyloid aggregate diversity, leading to the formation of novel conformations and propagation behaviors. This highlights the potential risk associated with exogenous prion-like proteins or peptides derived from their proteolysis[48,49], as once they enter the body, they can indirectly promote the formation of additional, more toxic aggregated structures (Figure 1A). Therefore, observing no or minimal effects in a single-step aggregation assay may be misleading, as it does not consider the seeding capacity of newly formed aggregates, which can only be assessed through subsequent aggregation assays (Figure 1C). Taken together, our results support that the molecular interactions Aβ experiences during its journey through the body (referred to here as Interaction History, Figure 1C) may be critical factors for understanding the disease and developing effective therapeutic approaches.

## Results and Discussion

### Exogenous Prion-like Sequences Can Influence Aβ Peptides Aggregation

In our previous investigation of the gut microbiome, we build a collection of ten exogenous prion-like sequences capable of interfering with Aβ40 aggregation[47] (Figure 1B). These sequences were selected based on the amyloid stretch hypothesis, which states that specific regions within a protein, known as amyloid cores, can drive amyloid nucleation[50–52]. These ten prion-like sequences are 21-residue peptides with net charges ranging from +2 to −2 and compositional similarities below 30% (Table 1)[47]. Since these sequences originate from gut microbiome bacteria, they may enter the gut lumen either through active export or after bacterial cell lysis[48,49] (Figure 1A), becoming free to interact with host proteins throughout the body. Based on this premise, in the present study, we used this collection as a model to investigate how exogenous molecules influence the interplay between Aβ40 and Aβ42, with a particular focus on heterotypic interactions and cross-seeding events (Figure 1C).

**Table 1.** Set of Exogenous Prion-like Sequences. Columns (from left to right) display: sequence identifiers used in this study; UniProt accession codes; source species from which the sequences originate; amino acid sequences, with residues colored according to the Zappo color scheme, aliphatic/hydrophobic (ILVAM) - rose, aromatic (FWY) - orange, positive (KRH) - dark blue, negative (DE) - red, hydrophilic (STNQ) - light green, conformationally special (PG) - magenta, cysteine (C) - yellow. Sequences obtained from Curto, J.S. et al.[47]. Net charge measured with Protein Calculator V3.4.

| Name | Uniprot ID | Species | Sequences | Net charge |
| --- | --- | --- | --- | --- |
| HP1 | M5YJZ4 | *Helicobacter pylori* | G N N N N S V I S F N Q T N F N Q G T Y N | -0,1 |
| HP2 | M3SJI9 | *Helicobacter pylori* | S S V Y W L N S V N E N N N N K S Y Y I S | -0,1 |
| CC3 | C0B555 | *Coprococcus comes* | M Q A A F N F I N N R R Y Q D A I N V L N | 0,9 |
| BH4 | C9L6N5 | *Blautia hansenii* | Y Q Q A L N V L S R I Q N R N A Q W Y Y L | 1,9 |
| RI5 | D4KZ46 | *Roseburia intestinalis* | H A N E G N V N Q Y N N S Y Q N T N S Y S | -0,9 |
| RI6 | C0FUS6 | *Roseburia inulinivorans* | M D N N Y Y N N N N N N N T N S N S N Y N | -1,1 |
| SA7 | E7GXS5 | *Streptococcus anginosus* | R S G N Y V F L A N S T N R Y Q G N Y Y N | 1,9 |
| LD8 | F0HUU2 | *Lactobacillus delbrueckii* | S Y S S S A N N Y G Y G S N Y N Y S Y S A | -0,1 |
| P9 | F5LFB6 | *Paenibacillus* | G Y V Q S S Y G Q N Q F Q N S Q Y G S Y Q | -0,1 |
| HA10 | G9Y7N7 | *Hafnia alvei* | S T G Y S D L N S T Y N Q S A L I N Q I G | -1,1 |

To mimic the complex aggregation processes observed in living organisms, we designed an approach based on sequential heterotypic seeding assays (Figures 1A and 1C). In this study, we incorporate the interplay between Aβ42 and Aβ40, hypothesizing that the higher abundance of Aβ40[4,6] facilitates encounters with exogenous molecules, thereby indirectly affecting Aβ42 aggregation and disease progression. To explore this interplay, we monitored Thioflavin-T (Th-T) fluorescence (Figures 2-4) and collected transmission electron microscopy (TEM) images (Supplementary Figures 1 and 2) to assess how Aβ42 aggregation is influenced by pre-aggregated samples assembled under different heterotypic interactions. Through these assays, we aim to shed light on the possible scenarios Aβ42 might encounter as it moves through the body (Figure 1A) and elucidate how the “Interaction History” could impact its aggregation and propagation (Figure 1C).

In our previous study[47], we found that most of the bacterial prion-like sequences (8 out of 10) accelerated Aβ40 aggregation, with lag time correlating with the sequences’ net charge. Hence, positively charged sequences led to faster aggregation, while negatively charged ones had little effect or even slowed aggregation[53–55]. These results highlight the role of electrostatic forces in intermolecular interactions and align with earlier reports showing that negatively charged peptides inhibit Aβ aggregation[53,54]. In the present study, we extended our analysis to Aβ42 by incubating it with these pre-aggregated bacterial peptides. Similar to the effects observed with Aβ40, incubation of Aβ42 with a 1:10 molar ratio of prion-like sequences predominantly enhances its aggregation (seven samples presented shorter half-time, Figure 2).

**Figure 2.**
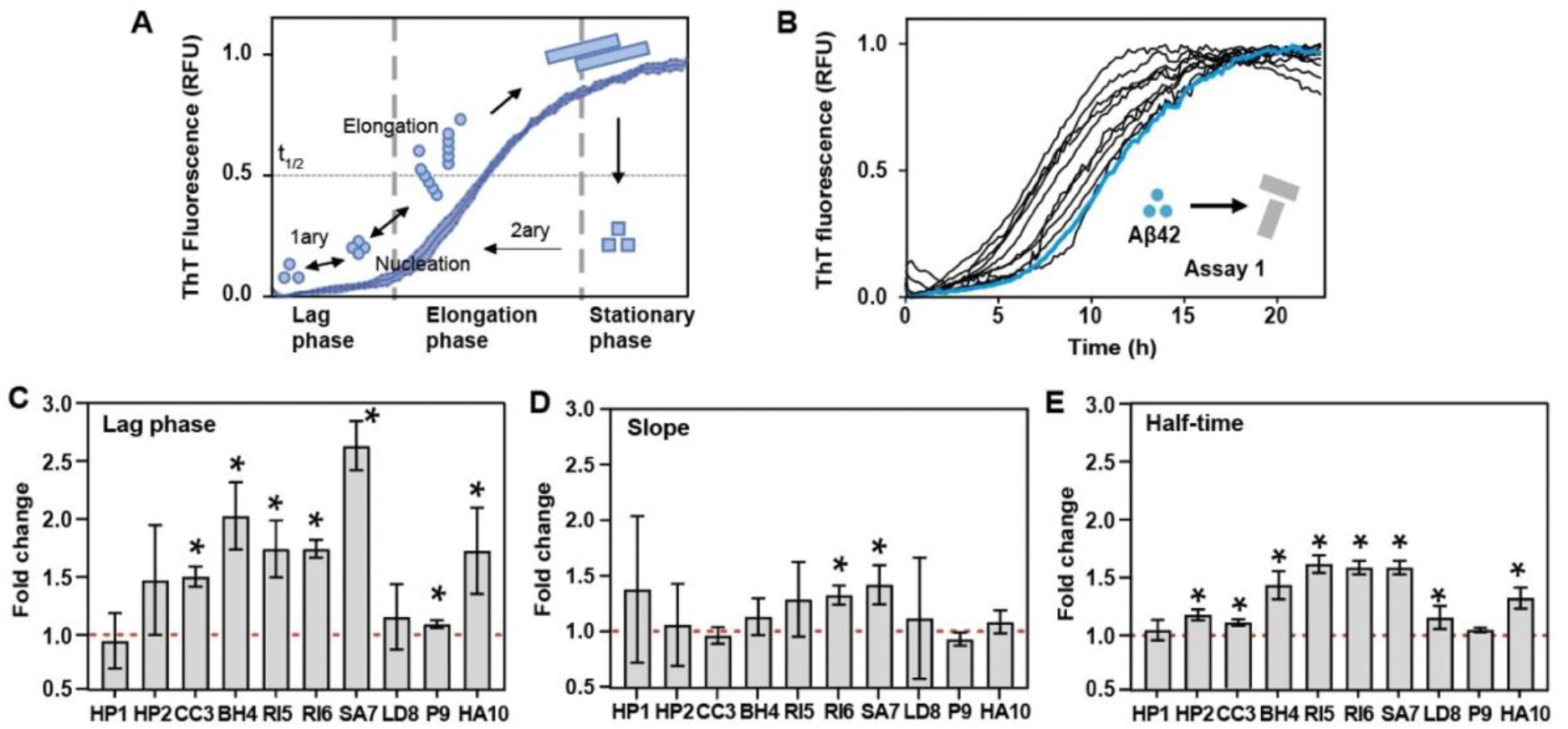
Exogenous Prion-like Sequences Influence Aβ42 Aggregation. A) Diagram showing the different phases of a protein aggregation process, indicating the different assembly mechanisms involved. B) Aggregation kinetics of Aβ42 in the presence (HP1–HA10, black lines) and absence (w/o, blue line) of seeds from exogenous prion-like sequences. See Supplementary Figure 4 for individual curves. C) Fold change in lag time (unseeded/seeded). D) Fold change in slope (seeded/unseeded). The equation fittings are shown in Figure 5. E) Fold change in half-time (unseeded/seeded). The red dashed line serves as a visual reference indicating a fold change of one relative to the aggregation assay started only with Aβ42 monomers (unseeded). An asterisk (*) indicates a statistically significant difference (p < 0.05) compared to the condition without seeds, as determined by an unpaired t-test. Error bars indicate the standard deviation. Each condition corresponds to the results of three biological replicates, with three technical replicates per biological replicate.

Although we observed only a correlation with the absolute net charge of bacterial prion-like sequences (Supplementary Figure 3), it is noteworthy that the peptides most strongly accelerating Aβ40 aggregation (SA7 and BH4)[47] also led to shorter lag times and half-times in Aβ42 aggregation kinetics (Figure 2C). In general, the observed enhancement in the aggregation curves appears to be primarily attributed to a shorter lag phase, as only two samples exhibited larger slopes, while seven displayed shortened lag times (Figure 2B-D). A reduced lag time indicates a more efficient primary nucleation phase, involving oligomeric species that serve as templates, whereas a steeper slope indicates a faster aggregation rate during the growth phase, which may result from the rapid addition of monomers to existing fibrils and could also involve secondary nucleation events (Figure 2A). The half-time (t1/2) serves as an indicator of overall aggregation efficiency, influenced by both lag phase and slope. Together, these findings suggest that these exogenous prion-like sequences primarily enhance the nucleation of Aβ42 aggregation by acting as seeding points (Figure 2).

**Figure 3.**
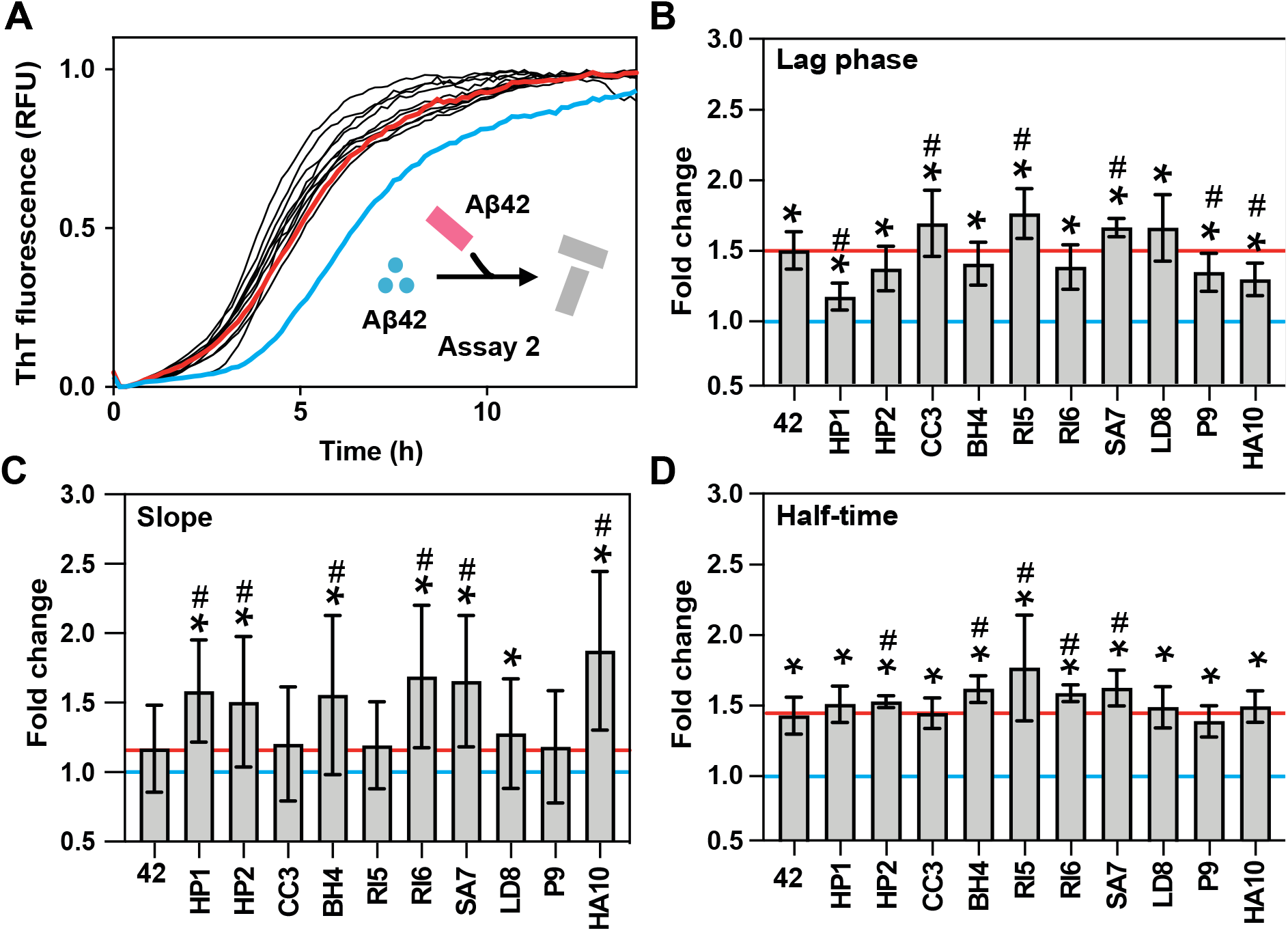
Exogenous Prion-like Sequences Affect Aβ42 Seeds and Their Propagation. A) Aggregation kinetics of Aβ42 seeded with Aβ42 aggregates formed in the presence and absence (w/o) of seeds from exogenous prion-like sequences. In blue and red are highlighted the aggregation kinetics of Aβ42 without seeds and with seed of pure Aβ42 aggregates, respectively. See Supplementary Figure 6 for individual curves. B) Fold change in lag time (unseeded/seeded). C) Fold change in slope (seeded/unseeded). The equation fittings are shown in Supplementary Figure 7. D) Fold change in half-time (unseeded/seeded). Horizontal blue line indicates one-fold change, the value corresponding to a kinetic of Aβ42 without seeds. Horizontal red line indicates the average value of the 42 column, the aggregation kinetics of Aβ42 seeded with pure Aβ42 aggregates. Error bars indicate the standard deviation. Each condition represents the results of three biological replicates, with three technical replicates per biological replicate. An asterisk (*) indicates a statistically significant difference (p < 0.05) compared to the condition without seeds, as determined by an unpaired t-test. A hash symbol (#) indicates a statistically significant difference (p < 0.05) compared to the condition seeded with pure Aβ42 aggregates, determined by an unpaired t-test.

### Effect on Aβ42 Seeds and Their Propagation

After observing that bacterial prion-like sequences influence the aggregation of both Aβ40[47] and Aβ42, we investigated how past molecular interactions affect the propagation of Aβ42 aggregates (Figure 1C). To address this, we conducted kinetic assays using pre-formed Aβ fibrils previously seeded with the bacterial peptides (Figure 2, Supplementary Figure 4). This second aggregation step (Figure 1C; Figure 3) involved a tenfold dilution of the exogenous peptides compared to the initial direct seeding assays. This dilution was achieved by incorporating 10% of the Aβ aggregates formed during the first step (Methods), keeping minimal the influence from the bacterial molecules (≈1%) and allowing us to focus on the properties of the pre-formed aggregates.

All lag times were accelerated compared to Aβ42 without seeds (Figure 3B, * symbol), with three bacterial sequences (CC3, RI5, SA7) even outperforming the kinetics seeded with pure Aβ42 aggregates, which were formed just by Aβ42 molecules (Figure 3B, # symbol). Regarding the slope, seven sequences exhibited faster elongation/secondary nucleation compared to unseeded monomers, while six out of ten presented significantly larger slopes than those triggered by the pure Aβ42 seeds. These findings point to an amplification effect, where the presence of bacterial peptides alter the Aβ42 aggregates, generating polymorphs with enhanced seeding capabilities. This also underscores the importance of the Interaction History, demonstrating that even exogenous molecules with limited initial seeding capacity can redirect the aggregation pathway toward a new population of aggregates with enhanced propagation potential. Additionally, while the first-step assay, in which bacterial sequences acted as seeds, principally influenced the primary nucleation phase, the subsequent aggregation step, seeded with the resulting Aβ42 aggregates, presented both a shorter lag time and a stepper slope, supporting the concept that sequence similarity favors the cross-seeding efficiency[56–58].

Beyond the overall aggregation acceleration, our analysis of different Aβ42 seeds revealed that each bacterial prion-like sequence produces fibrillar polymorphs with distinct seeding properties (Figure 3A-D, Supplementary Figure 6). Specifically, some bacterial peptides (SA7) reduce both lag time and slope, while others selectively influence only one parameter (HP1, HP2, CC3, BH4, RI5, RI6, HA10). Some sequences have no significant effects (LD8), whereas others even delay the aggregation process (P9). These findings indicate that the fibrillar polymorphs formed in the presence of different exogenous molecules have distinct properties and can influence different phases of host protein aggregation. This, in turn, affects subsequent interactions and aggregate propagation, even during later stages when the exogenous molecule is no longer present. For instance, Aβ42 seeds formed in the presence of RI5 shorten lag phase, suggesting enhanced monomer adherence during the initial nucleation phase, whereas Aβ42 seeds formed in the presence of BH4 increase the slope, potentially indicating accelerated elongation driven by secondary nucleation events. This diversity in aggregation-modulating mechanisms may contribute to increase the structural heterogeneity of amyloid fibrils, further complicating the pathological landscape of AD.

**Figure 4.**
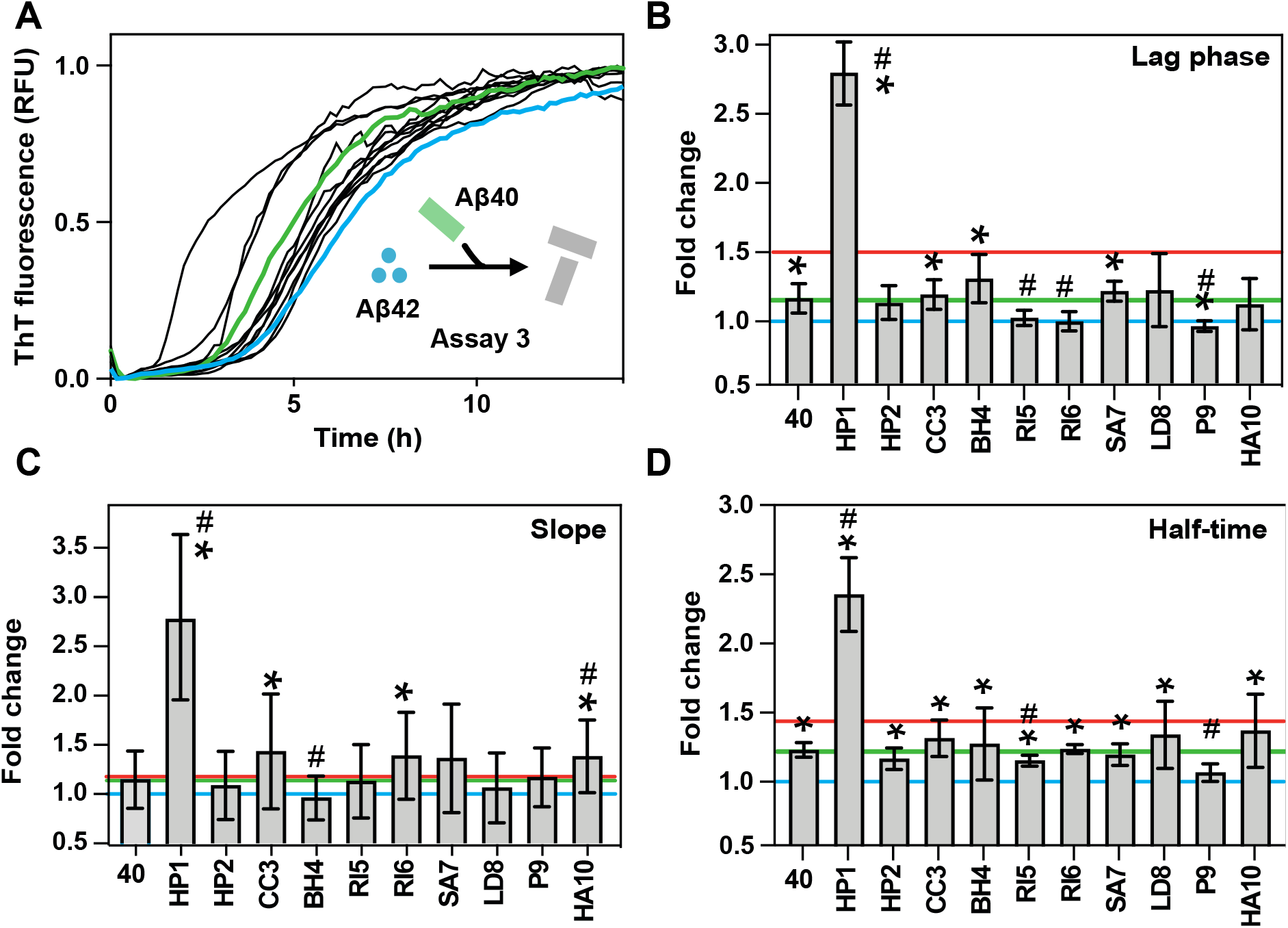
Exogenous Prion-like Sequences Affect Aβ40 Aggregation and Its Propagation to Aβ42. A) Aggregation kinetics of Aβ42 seeded with Aβ40 aggregates formed in the presence (HP1–HA10) and absence (w/o) of seeds from exogenous prion-like sequences. See Supplementary Figure 8 for individual curves. Green and blue lines highlight the aggregation of Aβ42 seeded with pure Aβ40 aggregates or without seeds, respectively. B) Fold change in lag time (unseeded/seeded). C) Fold change in slope (seeded/unseeded). The equation fittings are shown in Figure 9. D) Fold change in half-time (unseeded/seeded). The horizontal lines indicate the average value for aggregation kinetics of Aβ42 seeded with pure Aβ42 aggregates (red), pure Aβ40 aggregates (green) or without seeds (blue). Error bars indicate the standard deviation. Each condition represents the results of three biological replicates, with three technical replicates per biological replicate. An asterisk (*) indicates a statistically significant difference (p < 0.05) compared to the condition without seeds, as determined by unpaired t-test. A hash symbol (#) indicates a statistically significant difference (p < 0.05) compared to the condition seeded with pure Aβ40 aggregate, determined by an unpaired t-test. Blue Line represents the aggregation kinetics of Aβ42 without any seeds.

### Effect on Aβ40 and Its Subsequent Seeding of Aβ42

While the previous experiments focused on the self-seeding of Aβ42, it is important to consider that Aβ40 is the predominant isoform of amyloid-beta peptide in the human body[4,6]. Given its higher abundance, Aβ40 is more likely to encounter and interact with exogenous prion-like proteins. To explore this possibility, we examined whether our collection of exogenous sequences could also influence the cross-seeding potential of Aβ40 on Aβ42 (Figure 4, Supplementary Figure 8).

In previous studies, we observed that most bacterial sequences, from our collection, accelerated the aggregation of monomeric Aβ40[47]. In the present study, we used Aβ40 aggregates formed during this initial aggregation step to seed Aβ42 deposition. As a control, we tested the seeding efficiency of pure Aβ40 aggregates and found that, although they reduced the lag time and half-time of Aβ42 aggregation to a lesser extent than Aβ42 seeds (Figure 4B and D), they induced a comparable slope increase (Figure 4C). These findings support that residues 41 and 42 play a critical role in facilitating the assembly of free monomers during the primary nucleation process, and their absence in Aβ40 reduces its propagation efficiency[59].

When analyzing the effects of Aβ40 aggregates formed in the presence of the bacterial peptides, we observed that only four of the ten Aβ40 seeds shortened the lag phase, and among these, only one (HP1) achieved a lag time shorter than that of pure Aβ40 aggregates (Figure 4B). Interestingly, three of these Aβ40 seeds also enlarged the slope, validating that incubation with bacterial molecules can generate fibrillar polymorphs capable of influencing distinct aggregation phases. Overall, seven out of ten peptides redirected the aggregation pathway toward a new population of aggregates that significantly reduced the half-time of Aβ42 aggregation (Figure 4D). However, only in presence of HP1, Aβ40 seeds achieve a half-time reduction shorter than that of pure Aβ40, and more notably, even shorter than pure Aβ42 aggregates (Figure 4D, red line, p-value < 0.05, unpaired t-test). In contrast, some bacterial peptides, such as RI5 and P9, generated Aβ40 aggregates that slowed the Aβ42 aggregation (Figure 4, Supplementary Figure 8).

**Figure 5.**
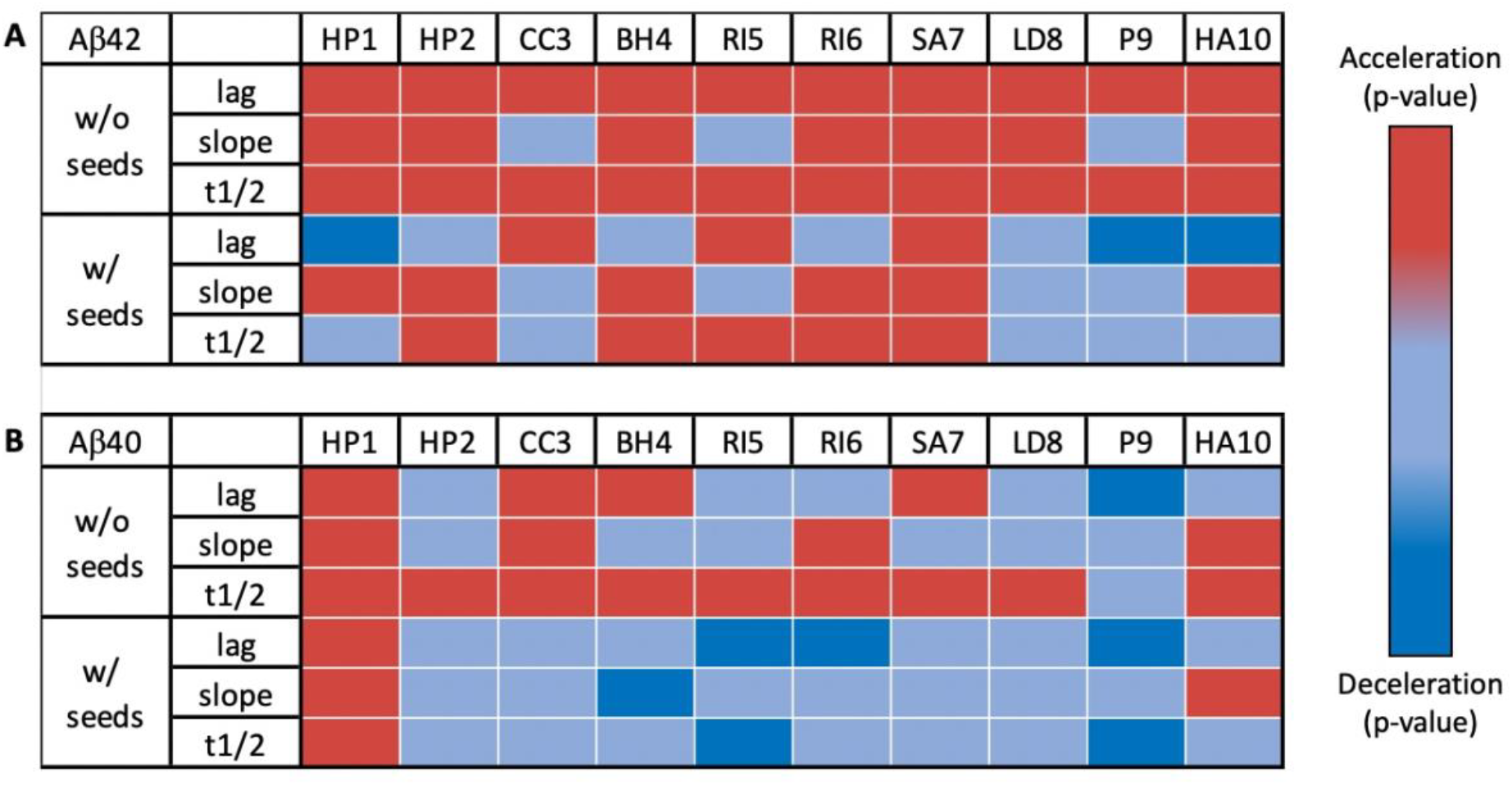
Differential Effects of Exogenous Prion-like Sequences on Aβ42 Aggregation. This figure provides a schematic overview of the significant data presented in Figures 3 and 4 (asterisk and hash symbols). It includes two color-coded matrices illustrating the impact of exogenous prion-like sequences on Aβ42 aggregation when occurring as a secondary or posterior seeding event. Red indicates conditions that significantly accelerate Aβ42 aggregation. Light blue represents conditions with no significant changes. Dark blue indicates conditions that significantly decelerate Aβ42 aggregation. Panel A show the results from assays using Aβ42 aggregates as seeds. Panel B show the results from assays using Aβ40 aggregates as seeds. The first row of each matrix displays the specific exogenous sequences added during seed formation. The first column indicates the statistical significance. w/o seeds refers to comparisons against the aggregation kinetics of Aβ42 without any seeds. w/ seeds refers to comparisons against Aβ42 aggregation when seeded with (A) pure Aβ42 aggregates or (B) pure Aβ40 aggregates. The second column lists the parameters analyzed, from top to bottom: Lag time, Slope and Half-time (t1/2).

The limited effect of Aβ40 aggregates, incubated with or without exogenous peptides, on Aβ42 aggregation (Figure 5) suggests that Aβ40 may have evolved to resist harmful interactions with exogenous molecules[60,61], making it less likely to propagate an aggregative conformation throughout the organism. Interestingly, HP1, the only sequence able to break this resilience, originates from a putative vacuolating cytotoxin of *Helicobacter pylori*, a bacterium associated with AD[62,63]. Consistent with this, our previous work[47] demonstrated that the HP1 sequence caused one of the most pronounced negative effects on *C. elegans* cognitive abilities.

Our findings highlight the potential of exogenous amyloid sequences to influence both direct seeding and subsequent cross-seeding events between Aβ isoforms. This effect varies among exogenous molecules, suggesting that specific molecular properties determine both the extent and direction of their influence, a feature that could inspire novel therapeutic strategies. Moreover, the structural heterogeneity of amyloid deposits, as evidenced by multiple fibril polymorphs identified in patient brain samples[10,25,44,64], suggests that exogenous prion-like sequences may further diversify aggregate populations, adding complexity to the pathological landscape of AD.

**Figure 6.**
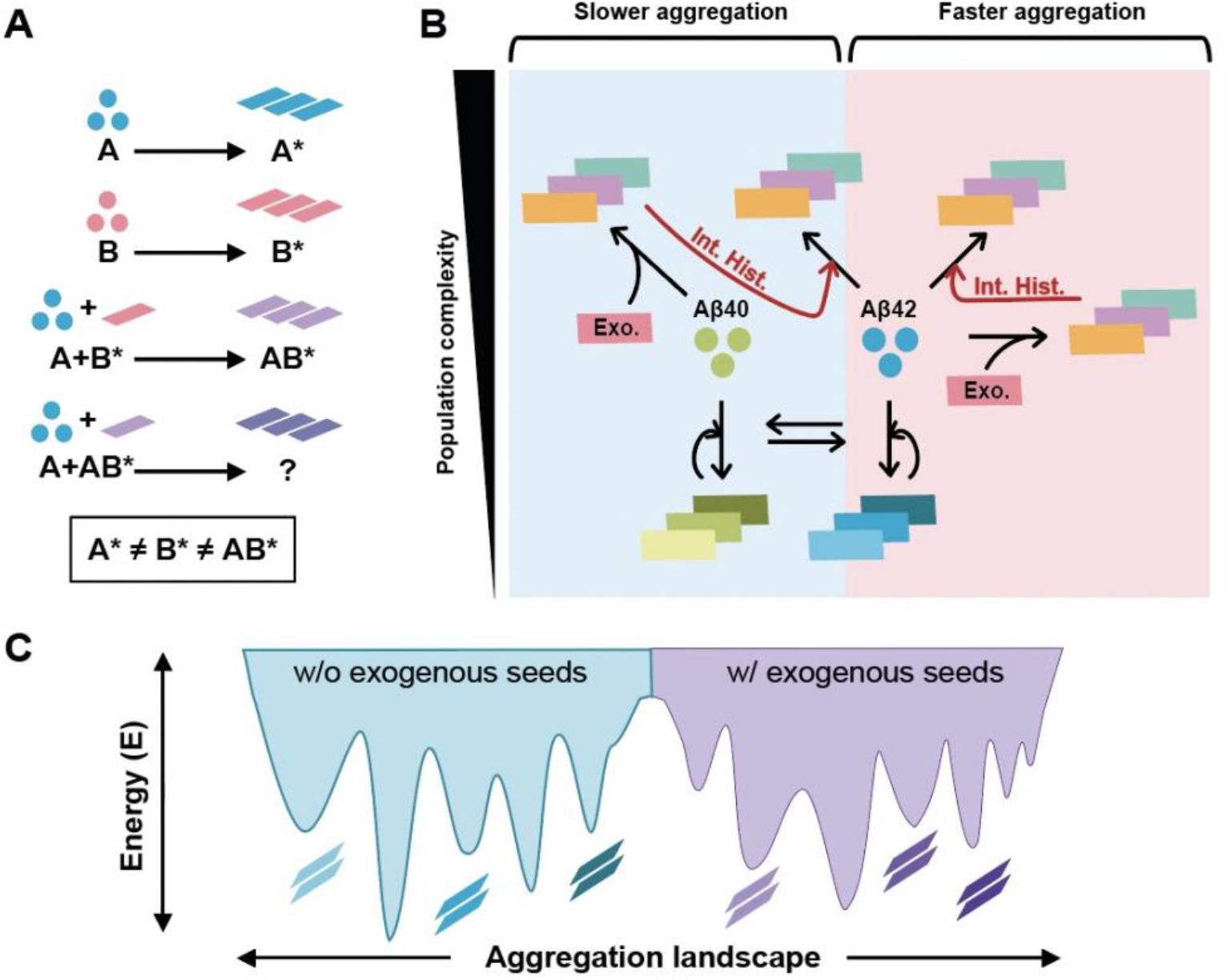
Influence of exogenous molecules on Aβ aggregate formation. A) This schematic illustrates how an exogenous protein (B) influences the aggregation dynamics of a host protein (A). Top row: The host protein (A) self-aggregates into fibrils (A*). Second row: The exogenous protein (B) also undergoes self-aggregation, forming fibrils (B*). Third row: When monomeric A aggregates in the presence of preformed B* fibrils, a distinct aggregate (AB*) is formed, different from A* or B*. Fourth row: A subsequent seeding step, where monomeric A interacts with AB* fibrils, results in an unknown aggregation outcome, suggesting that the aggregation pathway is shaped by prior molecular interactions, leading to fibrillar polymorphism. Bottom statement: The divergence between A, B, and AB* indicates that fibril formation is not uniform, which highlights the context-dependent nature of protein aggregation, where prior molecular interactions shape the polymorphic spectrum of aggregates. B) The aggregation of Aβ40 and Aβ42 is a dynamic process influenced by multiple interactions occurring in parallel. The presence of exogenous amyloid-like sequences (pink rectangles) introduces additional variability, modulating aggregation kinetics and the structural properties of the resulting fibrils. Since aggregates can act as seeds in consecutive cross-seeding events (curved arrows), Interaction History (red arrows) determines the evolution of amyloid species. As a result, aggregate diversity increases compared to a scenario where aggregation occurs in isolation, without exogenous influences (bottom part). This model suggests that the aggregation landscape is highly heterogeneous and sensitive to environmental factors. Circles represent monomeric proteins, while rectangles represent amyloid fibrils. C) In the absence of exogenous seeds, a protein populates a defined aggregation landscape (blue area), characterized by a specific set of aggregated structures. The presence of exogenous seeds may alter this landscape (purple area), potentially expanding the range of possible aggregation states and leading to additional polymorphs.

### Conclusions

Our findings support that amyloid propagation is not merely a linear extension of existing fibrils, but a dynamic process shaped by prior molecular interactions (Interaction History). Each aggregation cycle generates structurally distinct populations[10,34,35,46] that differ in composition, morphology, and seeding capacity, even when originating from the same monomeric protein (Figure 6). While the same molecular mechanisms control both primary and secondary seeding events, aggregation conditions (e.g., pH, ionic strength, co-aggregating species) define the resulting fibrillar polymorphs[10,34,35,46]. This context dependency may contribute to the strain-like behavior of amyloid diseases[36,37], allowing distinct fibril species to emerge and selectively propagate under specific aggregation conditions.

The energy landscape of amyloid aggregation consists of multiple local minima, each corresponding to distinct fibrillar polymorphs[10,34,35,46]. The introduction of exogenous amyloid fibrils can reshape this landscape (Figure 6C), amplifying structural heterogeneity and influencing downstream aggregation events[34,35,46]. These findings highlight why single-step aggregation assays may be insufficient, since aggregates formed under different conditions generate seeds with distinct propagation capacities, influencing subsequent aggregation in unexpected ways. In our experiments, exogenous amyloid sequences led to the formation of Aβ42 fibrillar polymorphs[25,44,64] with unique propagation properties. This underscores the role of environmental factors, particularly amyloids from the gut microbiota[39,40,45], in modulating Aβ aggregation and potentially contributing to neurodegeneration[47]. Importantly, our results indicate that heterologous assemblies may amplify aggregation, accelerating fibril formation and potentially exacerbating disease progression. Additionally, since exogenous molecules can shape subsequent interactions, aggregate propagation may occur far from the original site of exposure, even in the absence of the initiating agent.

While this study provides new insights into how exogenous amyloid sequences influence aggregation dynamics, several intriguing questions remain open for further study. A deeper structural characterization of the resulting polymorphs is needed to clarify their specific differences and determine whether exogenous molecules are incorporated into fibrils, potentially contributing to their heterogeneity. Additionally, it remains unclear whether exogenous interactions expand or limit the diversity of amyloid polymorphs. Another key aspect to explore is the potential toxicity of these aggregates, as certain polymorphs may be more pathogenic than others. Beyond the plateau phase, it is also essential to determine whether fibrillar polymorphs formed at different kinetic phases[34,35] exhibit distinct propagation properties and toxicity. This is particularly relevant in a biological system, where amyloid species from various aggregation stages may coexist, and any of them could potentially act as a trigger for acute disease onset.

Overall, this work emphasizes the need for a broader perspective on amyloid aggregation, one that considers external molecular influences, aggregation history, and fibril polymorphism. Understanding these variables could provide valuable insights into disease mechanisms and inspire the development of therapeutic strategies to address the structural complexity of amyloid deposits in AD and related conditions.

## Methods

### Protein alignment and net charge

The alignments between the 10 different amyloid-forming core candidates were obtained using the KAling method within the Unipro UGENE software v43.0 [65]. Unipro UGENE software v43.0 was also used for sequence representation.

Net charge was calculated with PROTEIN CALCULATOR v3.4 at pH 7 (https://protcalc.sourceforge.net/).

### Peptide Preparation

All samples were prepared in low protein-binding microcentrifuge tubes (ThermoFisher Scientific, Waltham, MA, USA). The bacterial amyloid sequences were obtained in lyophilized form from the Peptide Synthesis Facility, Department of Experimental and Health Sciences, Universitat Pompeu Fabra (UPF). Peptides were initially solubilized in 1,1,1,3,3,3-hexafluoroisopropanol (HFIP), then divided into aliquots and dried overnight in a fume hood at room temperature. For the amyloid aggregation, we employed the same conditions as in Curto, J.S. *et al*. 2024[47], in brief, the peptides were dissolved in DMSO to maintain monomeric states, followed by dilution in 50 mM phosphate buffer (PB), pH 7.4.

Synthetic Aβ40 and Aβ42 were purchased from GenScript (Rijswijk, Netherlands). Stock solutions were prepared by dissolving 1 mg of peptide in a final concentration of 250 μM in 20 mM sodium phosphate buffer with 0.04% NH_3_ and NaOH to achieve a pH of 11. The absence of aggregates was assessed using dynamic light scattering[47,66]. The peptide solution was then sonicated for 10 minutes (Fisherbrand Pittsburgh, PA, USA, FB15051) without sweep mode and stored at −80°C. For seeding experiments, the peptide was diluted at a 25 uM concentration with 20 mM phosphate buffer (PB), pH 7.4. Then, if necessary, the specific seeds were added to a final concentration of 2 uM and the mixture was left to aggregate at room temperature for 24 hours before their analysis.

### Thioflavin-T Binding and Aggregation Kinetics

Thioflavin-T (ThT) was dissolved in Milli-Q water to a concentration of 5 mM, filtered through a 0.2-μm filter, diluted to 0.5 mM, and stored at −20°C. ThT fluorescence for aggregation kinetics was measured every 10 minutes, measuring ThT fluorescence with a 440-nm excitation filter and a 480-nm emission filter on a plate reader (TECAN Infinite+ NANO) with bottom optics. Samples were prepared in flat-bottom, black, non-binding 96-well plates (Greiner Bio-One), with 100 μL of sample added per well. Each condition was measured in triplicate.

For seeding experiments, Aβ40 and Aβ42 peptide stocks at pH 11 were diluted to a final concentration of 25 μM in 20 mM sodium phosphate buffer with 100 mM NaCl. The corresponding pre-aggregated peptides were sonicated for 5 minutes before being added to a final concentration of 2.5 μM. Control conditions included samples without Aβ42 but with the peptide, samples without Aβ42, and samples without peptides, all prepared with equivalent volumes of the corresponding buffers. All samples and controls were prepared in triplicate. The pH in each well was measured at the beginning and end of the experiment to confirm stability throughout the aggregation process and to verify consistency across conditions. ThT was added to a final concentration of 20 μM. Aggregation reactions were conducted at 37°C without agitation, with four independent replicates. The t_1_/_2_, defined as the time to reach 50% of the final fluorescence intensity, was measured for each condition. The lag phase was calculated as the time to reach 10% of the final fluorescence intensity. Each condition was measured with three experimental replicates, each consisting of three technical replicates. Also, the data were fitted into a sigmoidal curve using the Hill function:

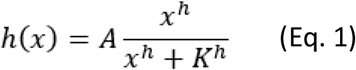

In short, A, h and K the fitting parameters. The fittings were used to calculate the slope of the exponential phase of the aggregation kinetic as done by Rupert et al[26].

### Transmission Electron Microscopy (TEM)

To prepare samples for TEM analysis, a 10 μL aliquot of the aggregated peptide solution (25 μM, incubated at 37°C for 24 hours without agitation) was applied to carbon-coated copper grids and allowed to adhere for 1 minute. Liquid excess was carefully blotted with Whatman filter paper[47]. Grids were then rinsed with distilled water and subsequently stained with 2% (w/v) uranyl acetate for 1 minute. After staining, grids were air-dried. Micrographs were obtained using a JEM-1400 transmission electron microscope (JEOL, Tokyo, Japan) set to an accelerating voltage of 80 keV.

### Statistics

GraphPad PRISM 8 software was used for statistical analysis of the obtained data. In general, the data was analyzed with by unpaired t-test. Statistical significance was considered with p-values <0.05. Data is shown as the mean ± SD.

## Acknowledgments

We extend our gratitude to the following individuals:

Jacob Rupert for his comments and in-depth discussions, which provided valuable insights into our work and its implications.

Transmission electron microscopy (TEM) images were acquired at the Servei de Microscòpia i Difracció de Raigs X (SMiDRX) of the Universitat Autònoma de Barcelona (UAB).

