## Supplementary Material for "Exogenous Amyloid Sequences: Their Role in Amyloid-Beta Heterotypic Aggregation"

Jofre Seira Curto[1]*, Marina Romero Ruiz[1], Genís Pérez Collell[1], Sandra Villegas[1], Maria Rosario Fernandez[1], Natalia Sanchez de Groot[1]*
[1]Unitat de Bioquímica. Departament de Bioquímica i Biologia Molecular. Universitat Autònoma de Barcelona, Spain.

**Supplementary Figure 1. TEM Images of Fibrillar Aggregates of Aβ42 Seeded with Aβ42.**

**Supplementary Figure 2. TEM Images of Fibrillar Aggregates of Aβ42 Seeded with Aβ40.**

**Supplementary Figure 3. Comparison between the net charge of peptides and their effect on Aβ42 aggregation kinetics.**

**Supplementary Figure 4. Aggregation kinetics of Aβ42 incubated with different bacterial amyloid cores.**

**Supplementary Figure 5. Aggregation kinetics of Aβ42 incubated with different bacterial amyloid cores, fitted to the Hill function.**

**Supplementary Figure 6. Aggregation kinetics of Aβ42 incubated with different seeds of Aβ42.**

**Supplementary Figure 7. Aggregation kinetics of Aβ42 incubated with different seeds of Aβ42, fitted to the Hill function.**

**Supplementary Figure 8. Aggregation kinetics of Aβ42 incubated with different seeds of Aβ40.**

**Supplementary Figure 9. Aggregation kinetics of Aβ42 incubated with different seeds of Aβ40, fitted to the Hill function.**

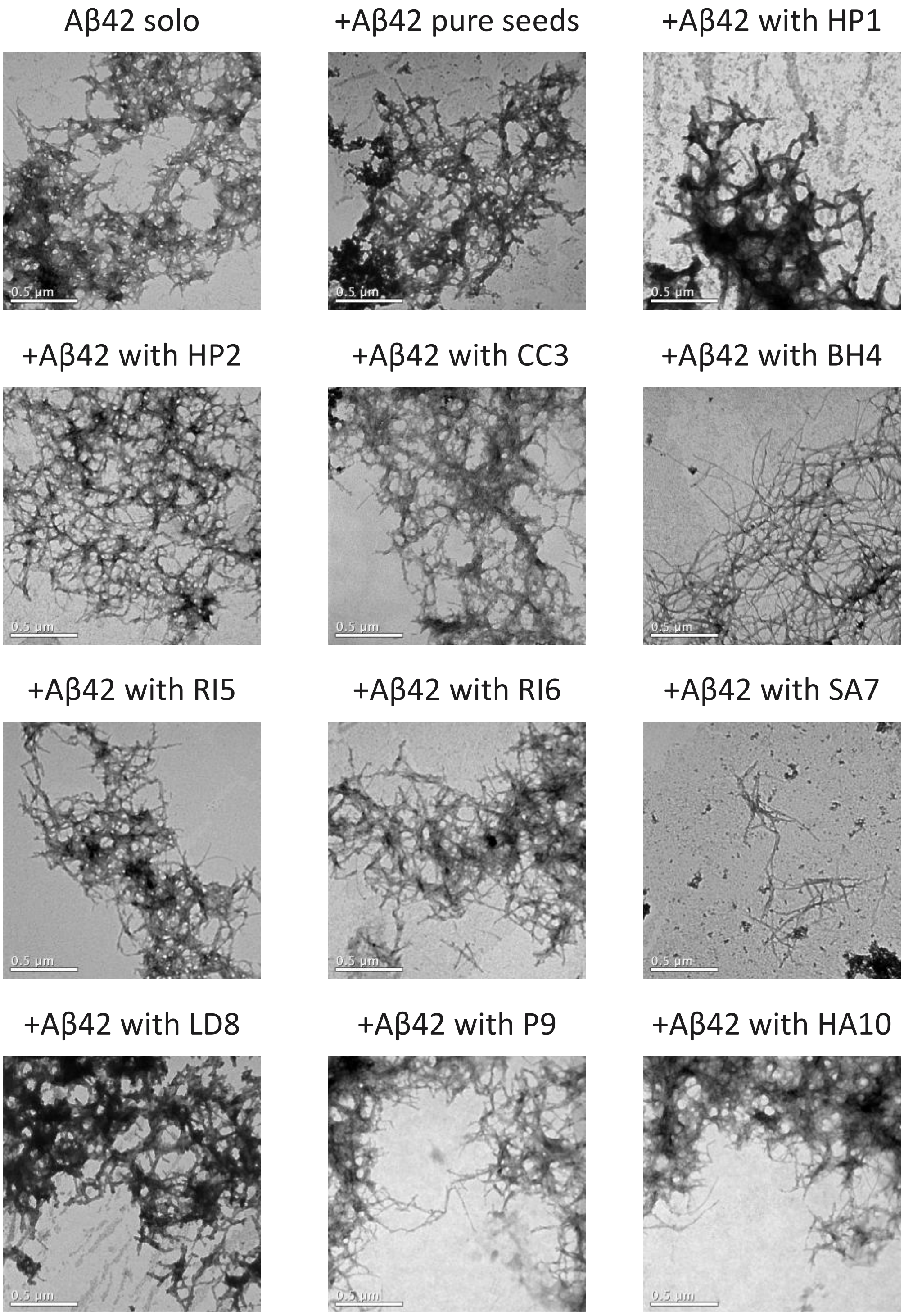


**Supplementary Figure 1. TEM Images of Fibrillar Aggregates of Aβ42 Seeded with Aβ42.** This figure presents transmission electron microscopy (TEM) images showing fibrillar aggregates of Aβ42 under different seeding conditions. The top left image shows Aβ42 aggregated without added seeds, while the top middle image shows Aβ42 aggregated with pure Aβ42 seeds. The remaining images illustrate Aβ42 aggregates seeded with Aβ42 fibrils formed in the presence of the indicated amyloid core sequences. All aggregates were formed after 24 hours of incubation at 37ºC without agitation.

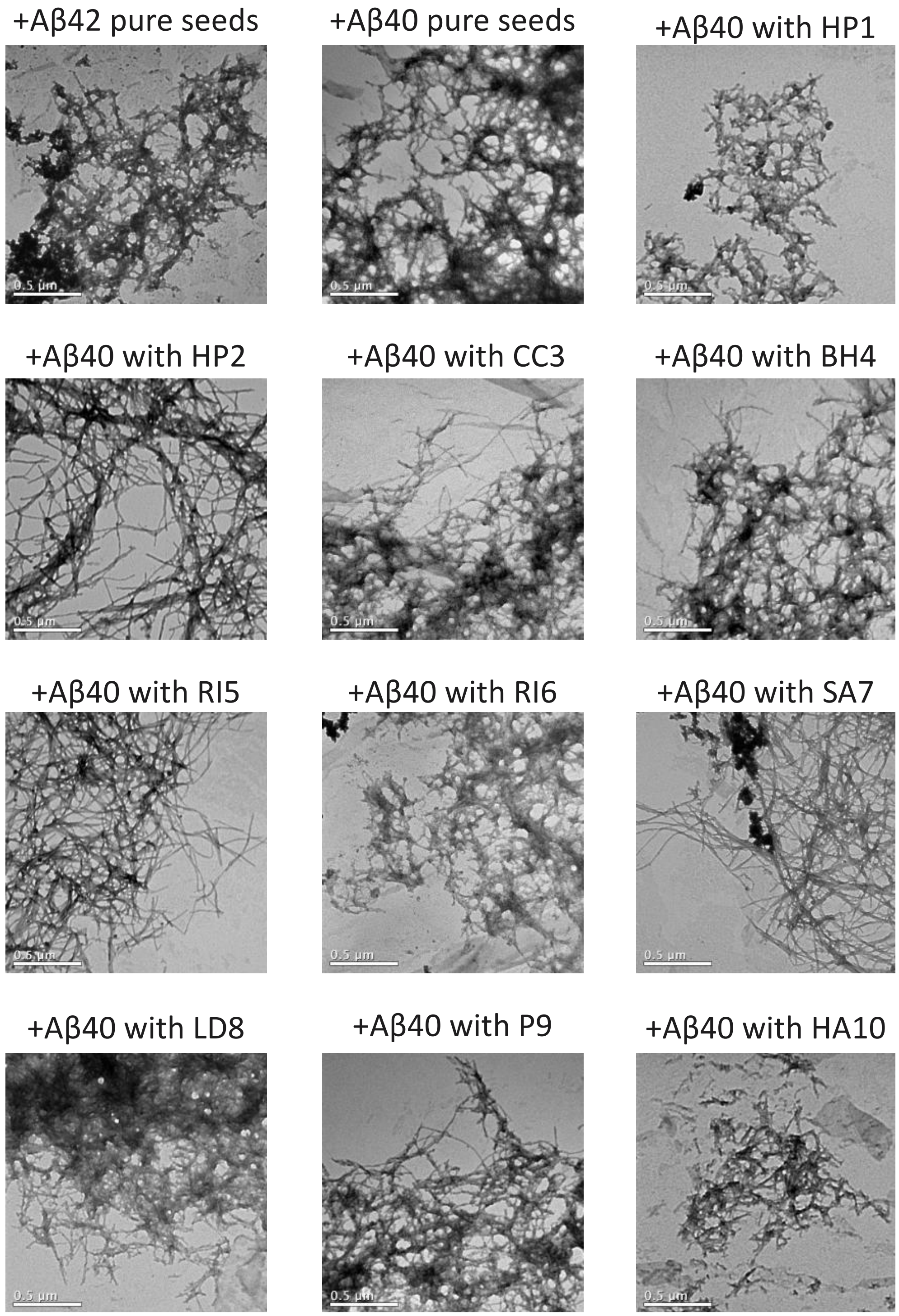


**Supplementary Figure 2. TEM Images of Fibrillar Aggregates of Aβ42 Seeded with Aβ40.** This figure presents transmission electron microscopy (TEM) images showing fibrillar aggregates of Aβ42 under different seeding conditions. The top left image shows Aβ42 aggregated with pure Aβ42 seeds, while the top middle image shows Aβ42 aggregated with pure Aβ40 seeds. The remaining images illustrate Aβ42 aggregates seeded with Aβ40 fibrils formed in the presence of the indicated amyloid core sequences. All aggregates were formed after 24 hours of incubation at 37ºC without agitation.

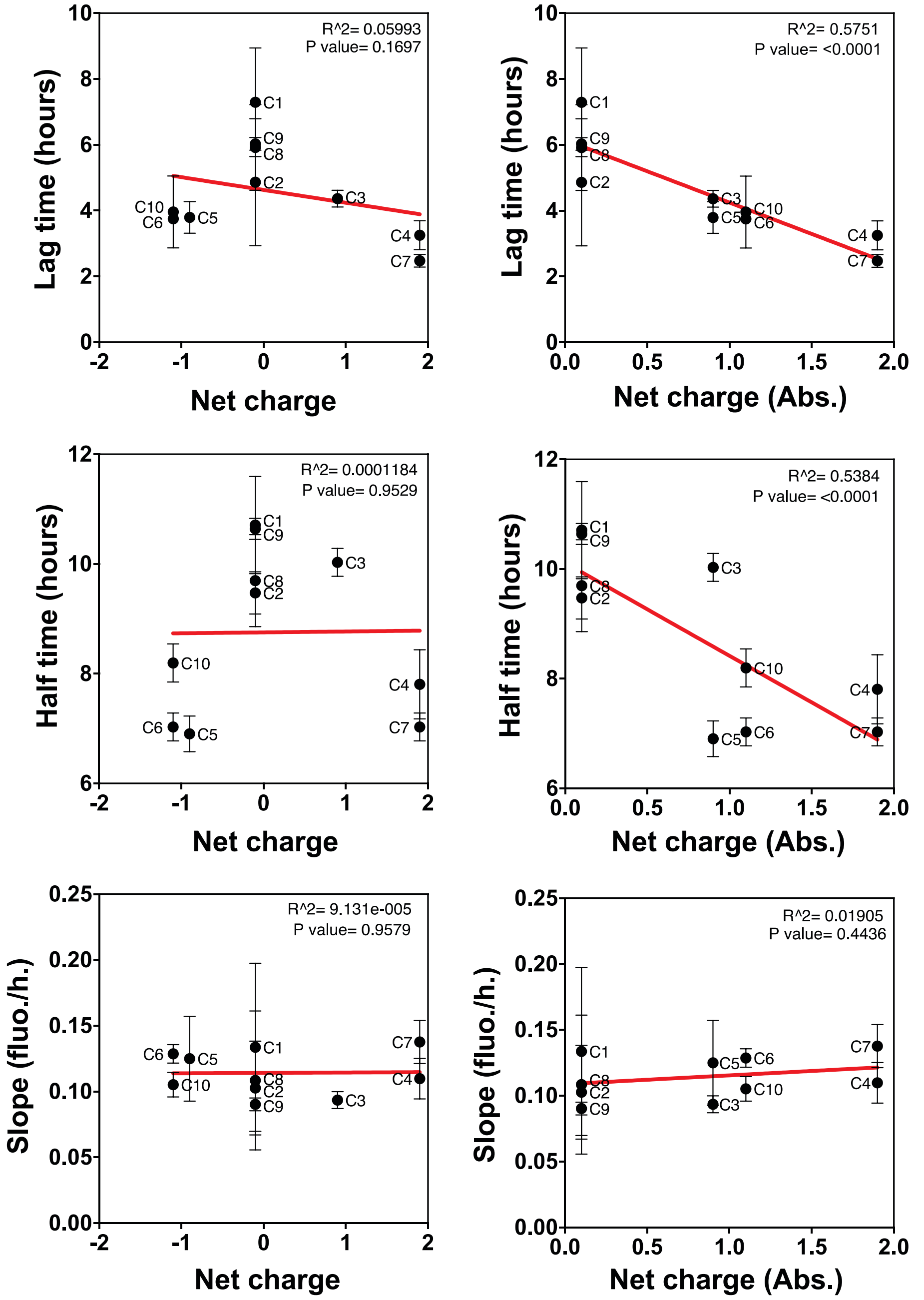


**Supplementary Figure 3. Comparison between the net charge of peptides and their effect on Aβ42 aggregation kinetics.** Each panel presents a scatter plot analyzing the relationship between peptide net charge (x-axis) and different kinetic parameters of Aβ42 aggregation (y-axis), including lag time (top row), half-time (middle row), and slope (bottom row). The left column considers the direct net charge values, while the right column evaluates the absolute net charge. The red lines represent linear regression fits, with $R^2$ values and p-values provided in each plot. Error bars indicate the standard deviation of the y-axis.

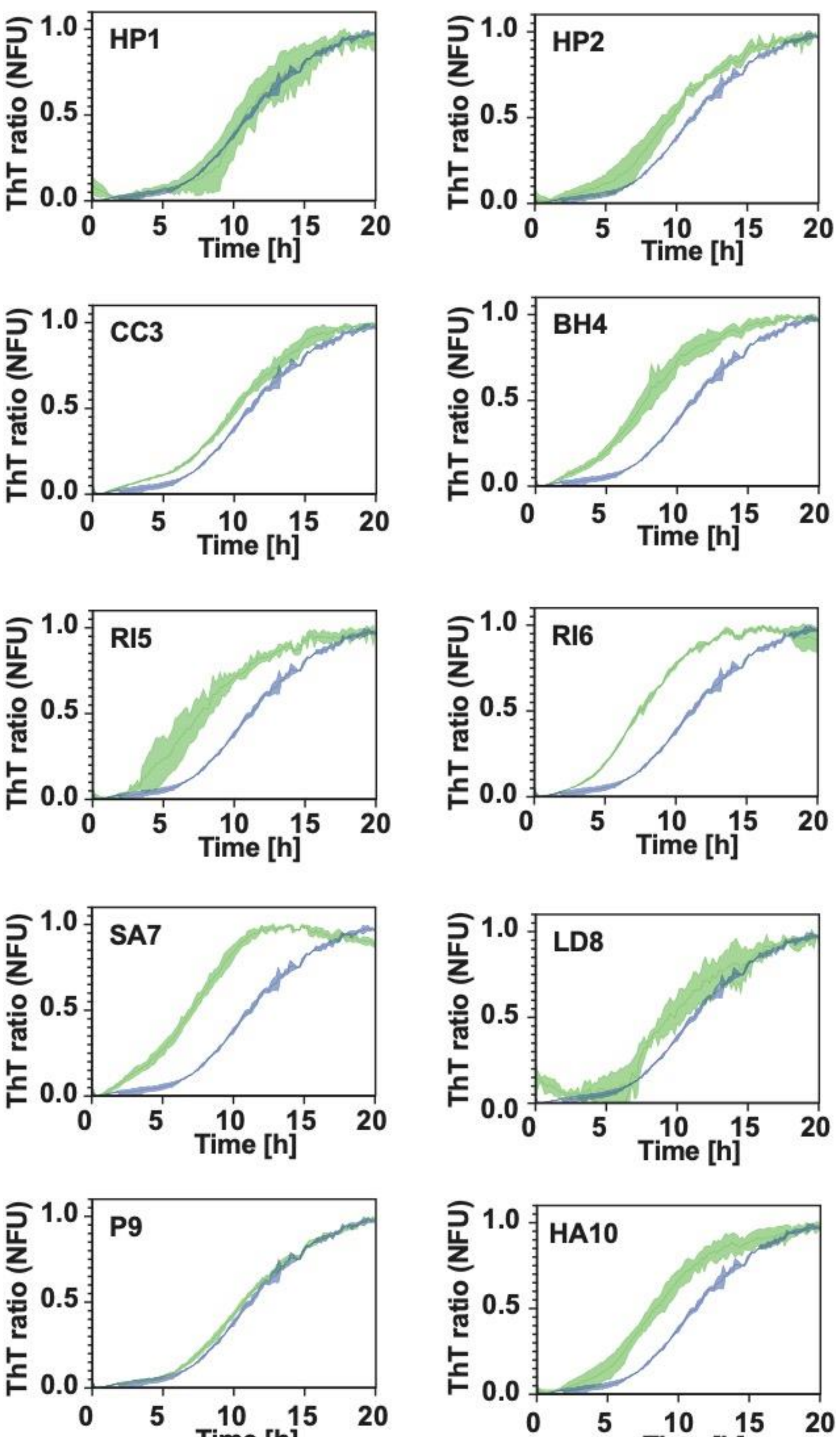


**Supplementary Figure 4. Aggregation kinetics of Aβ42 incubated with different bacterial amyloid cores.** The blue curve represents Aβ42 aggregation without seeds, while the green curve represents Aβ42 aggregation in the presence of pre-formed aggregates of the corresponding bacterial amyloid core. Each experiment was performed with three biological replicates, each containing three technical replicates.

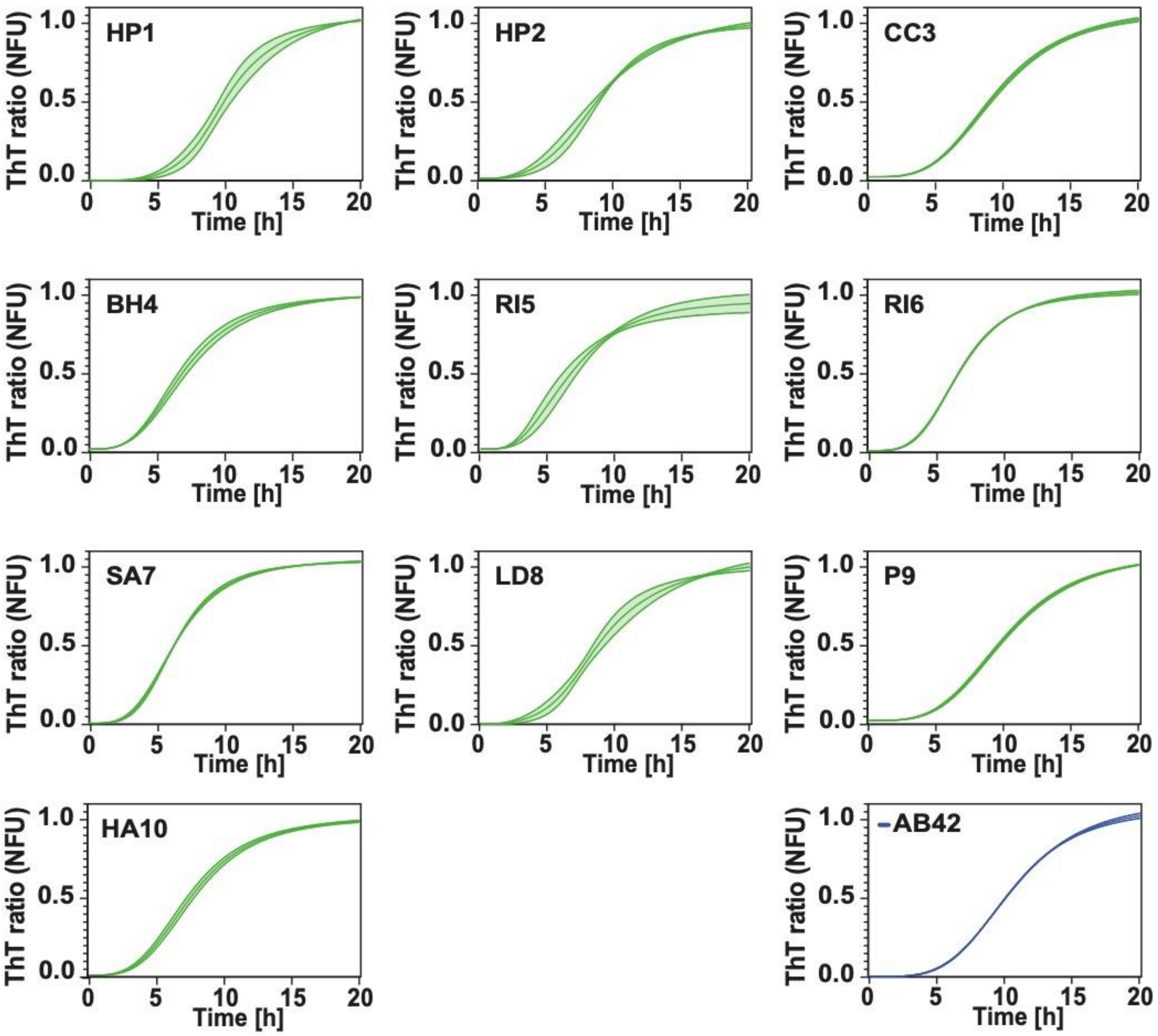


**Supplementary Figure 5. Aggregation kinetics of Aβ42 incubated with different bacterial amyloid cores, fitted to the Hill function.** The blue curve represents Aβ42 aggregation without seeds, while the green curve represents Aβ42 aggregation in the presence of pre-formed aggregates of the corresponding bacterial amyloid core. Each experiment was performed with three biological replicates, each containing three technical replicates. The middle line represents the fitted aggregation curve, and the shaded area corresponds to the standard deviation of the experimental replicates.

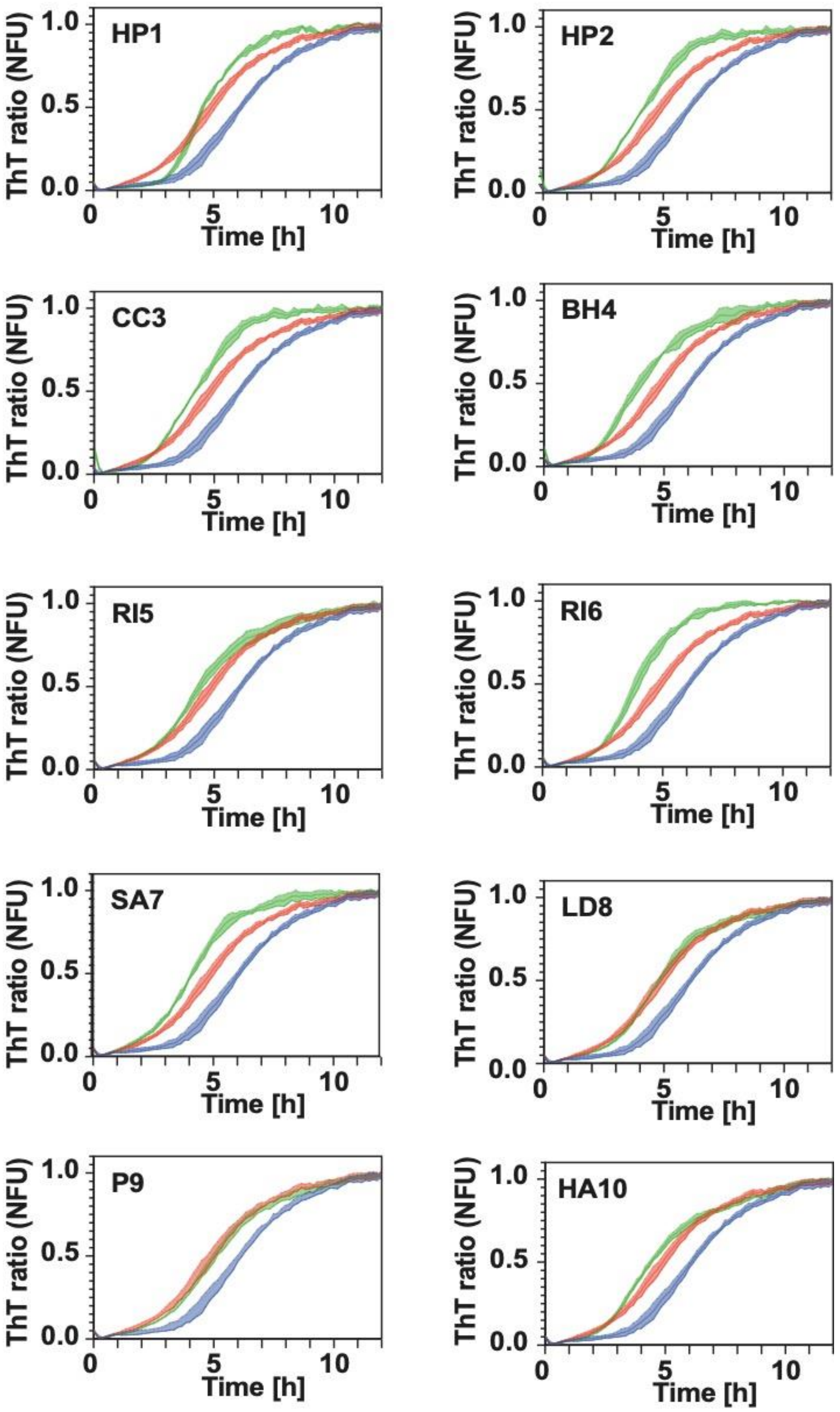


**Supplementary Figure 6. Aggregation kinetics of Aβ42 incubated with different seeds of Aβ42.** The blue curve represents Aβ42 aggregation without any seeding. The red curve represents Aβ42 aggregation in the presence of pure Aβ42 seeds. The green curve represents Aβ42 aggregation in the presence of Aβ42 seeds formed in the presence of the corresponding bacterial amyloid core. Each experiment was performed with three biological replicates, each containing three technical replicates.

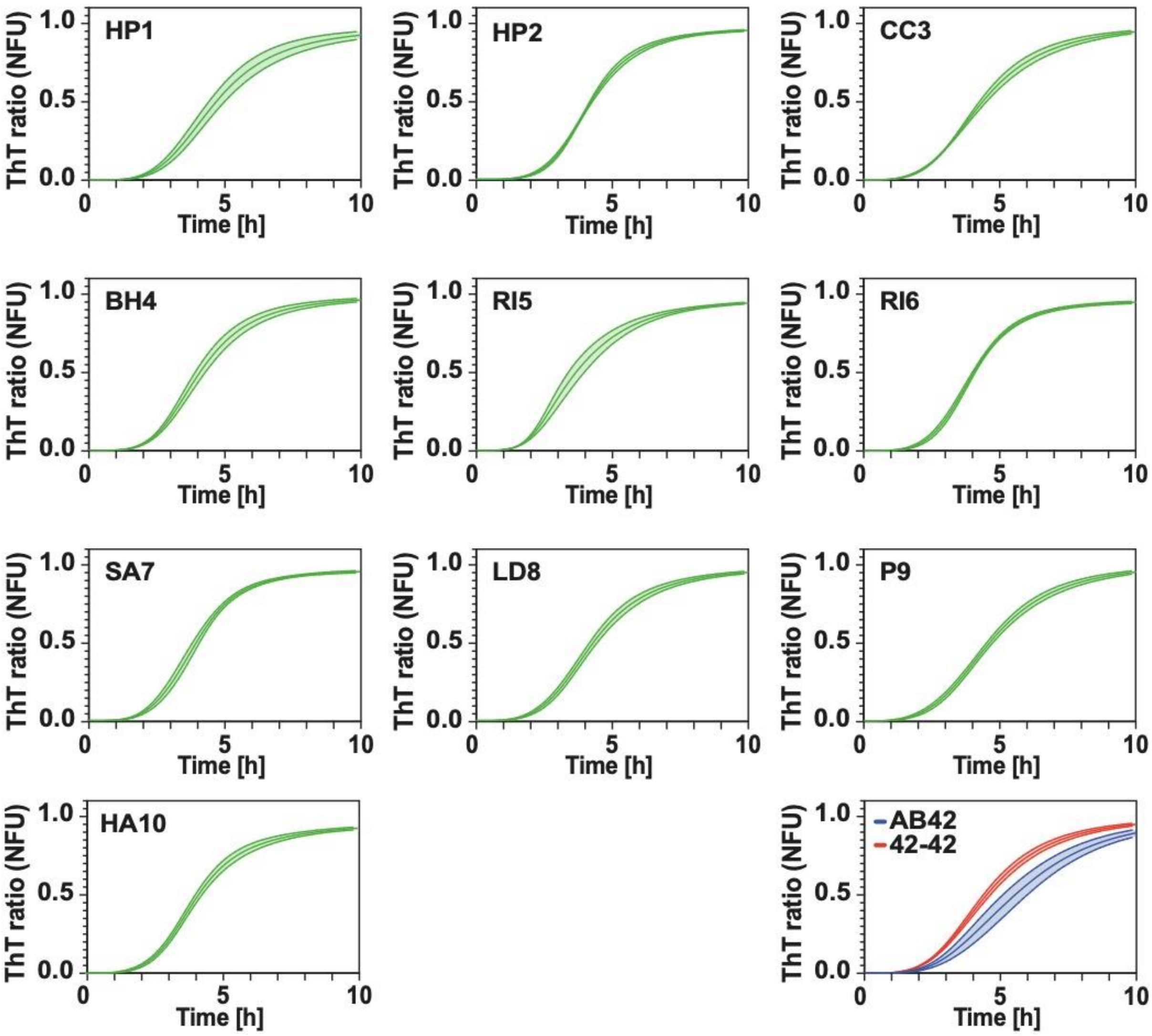


**Supplementary Figure 7. Aggregation kinetics of Aβ42 incubated with different seeds of Aβ42, fitted to the Hill function.** The blue curve represents Aβ42 aggregation without any seeding. The red curve represents Aβ42 aggregation in the presence of pure Aβ42 seeds. The green curve represents Aβ42 aggregation in the presence of Aβ42 seeds formed in the presence of the corresponding bacterial amyloid core. Each experiment was performed with three biological replicates, each containing three technical replicates. The middle line represents the fitted aggregation curve, and the shaded area corresponds to the standard deviation of the experimental replicates.

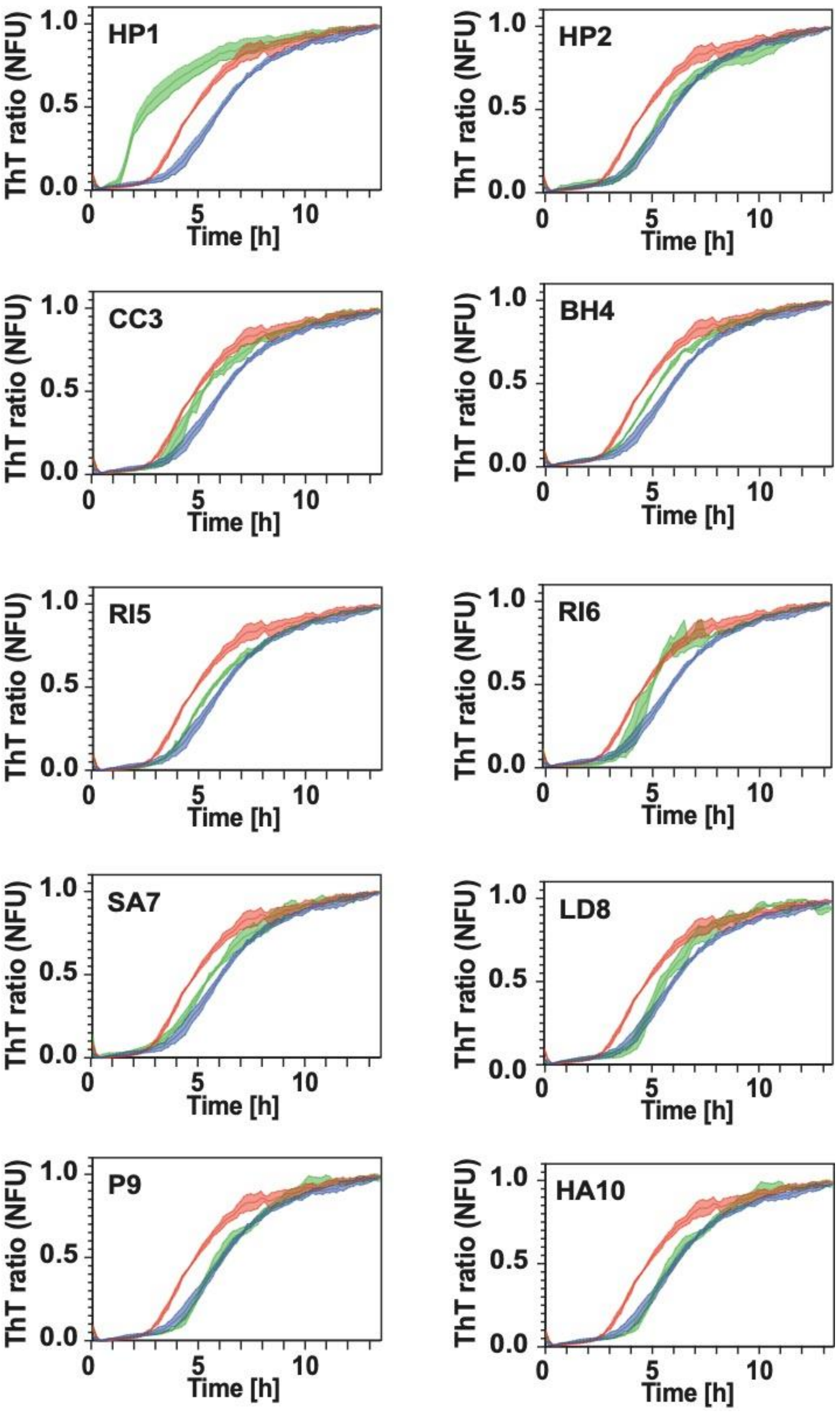


**Supplementary Figure 8. Aggregation kinetics of Aβ42 incubated with different seeds of Aβ40.** The blue curve represents Aβ42 aggregation without any seeding. The red curve represents Aβ42 aggregation in the presence of pure Aβ40 aggregates. The green curve represents Aβ42 aggregation in the presence of the Aβ40 aggregates formed in the presence of the corresponding bacterial amyloid core. Each experiment was performed with three biological replicates, each containing three technical replicates.

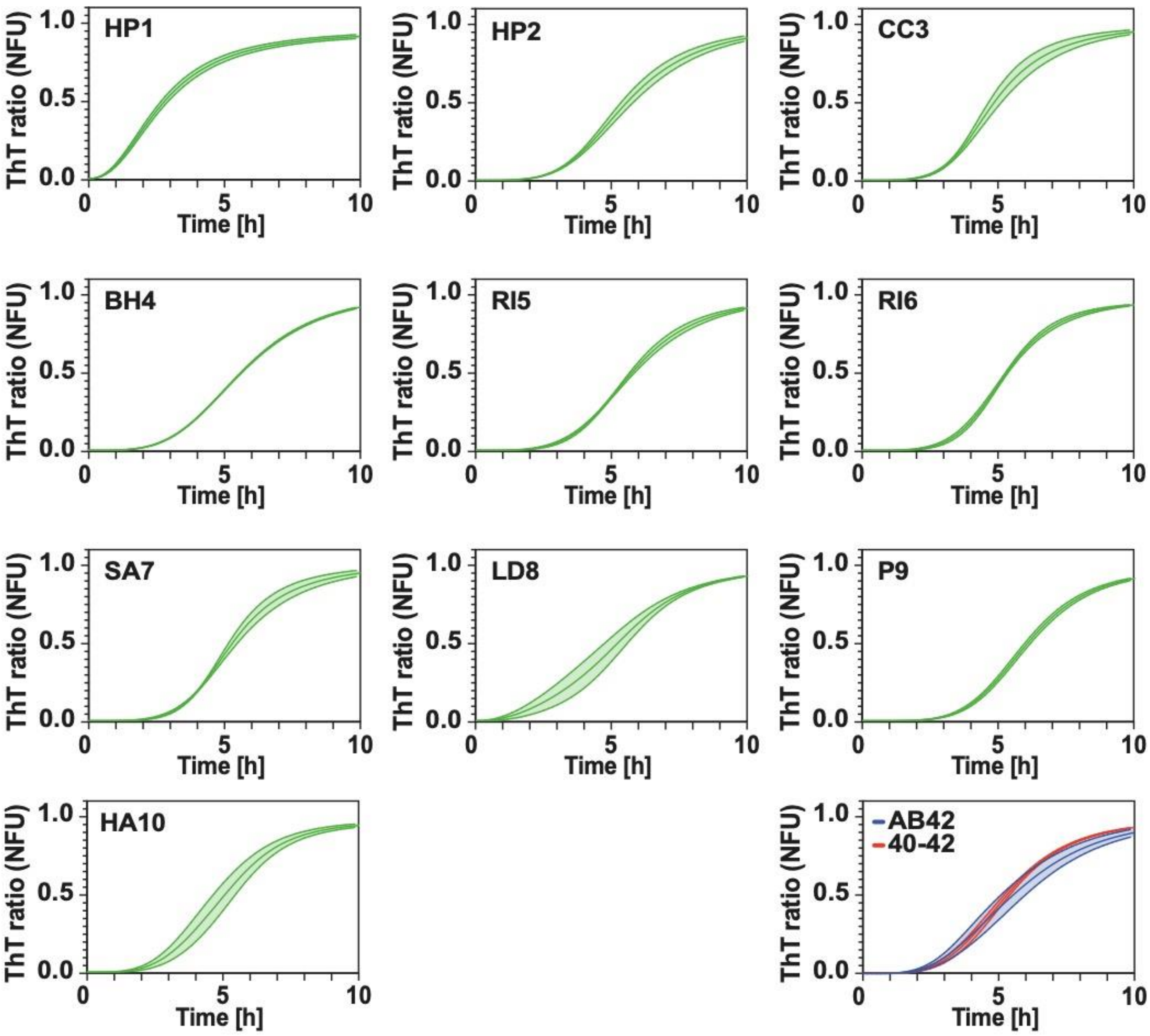


**Supplementary Figure 9. Aggregation kinetics of Aβ42 incubated with different seeds of Aβ40, fitted to the Hill function.** The blue curve represents Aβ42 aggregation without any seeding. The red curve represents Aβ42 aggregation in the presence of pure Aβ40 aggregates. The green curve represents Aβ42 aggregation in the presence of the Aβ40 aggregates formed in the presence of the corresponding bacterial amyloid core. Each experiment was performed with three biological replicates, each containing three technical replicates. The middle line represents the fitted aggregation curve, and the shaded area corresponds to the standard deviation of the experimental replicates.
